# Vitamin D induces SIRT1 activation through K610 deacetylation in colon cancer

**DOI:** 10.1101/2023.02.22.529558

**Authors:** José Manuel García-Martínez, Ana Chocarro-Calvo, Javier Martínez-Useros, María Jesús Fernández-Aceñero, M. Carmen Fiuza, Jose Cáceres-Rentero, Antonio De la Vieja, Antonio Barbáchano, Alberto Muñoz, María Jesús Larriba, Custodia García-Jiménez

## Abstract

Posttranslational modifications of epigenetic modifiers provide a flexible and timely mechanism for rapid adaptations to the dynamic environment of cancer cells. SIRT1 is an NAD^+^-dependent epigenetic modifier whose activity is classically associated with healthy aging and longevity, but its function in cancer is not well understood. Here, we reveal that 1α,25-dihydroxyvitamin D_3_ (1,25(OH)_2_D_3_, calcitriol), the active metabolite of vitamin D (VD), promotes SIRT1 activation through auto-deacetylation in human colon carcinoma cells, and identify lysine 610 as an essential driver of SIRT1 activity. Remarkably, our data show that the post-translational control of SIRT1 activity mediates the antiproliferative action of 1,25(OH)_2_D_3_. This effect is reproduced by the SIRT1 activator SRT1720, suggesting that SIRT1 activators may offer new therapeutic possibilities for colon cancer patients who are VD deficient or unresponsive. Moreover, this might be extrapolated to inflammation and other VD deficiency-associated and highly prevalent diseases in which SIRT1 plays a prominent role.

## INTRODUCTION

Over the last decades the study of genetic lesions in tumour cells provided key insights into the deregulated signalling pathways that promote cell proliferation and invasiveness and suppress senescence. In addition, increasing evidence from epidemiological studies suggests a significant impact of non-genetic modifiable factors on the oncogenic process by affecting cell epigenetics and thus, gene expression and phenotype. The key metabolic sensor SIRT1 acts an NAD^+^-dependent deacetylase and a critical post-translational modifier and epigenetic regulator. As such, SIRT1 promotes efficient energy utilisation and cellular defences in response to environmental challenges, whereas its dysregulation accelerates age-related diseases such as diabetes and cancer (Bonkowski and Sinclair, 2016).

Colorectal cancer (CRC) is the second most diagnosed malignancy in women, the third in men, and a leading cause of cancer-related deaths worldwide, with an incidence estimated to increase by 60% by 2030 (Bray et al., 2018; Cancer Genome Atlas Network, 2012). Many observational/epidemiological studies suggest that vitamin D (VD) deficiency is a risk factor for developing and dying of cancer, particularly for CRC (Feldman et al., 2014; Grant et al., 2022; Kim et al., 2022, 2021), albeit data from supplementation studies in the human population are controversial. Confirmation of a clinically relevant anti-CRC effect of VD in well-designed prospective randomized studies is still pending (Chandler et al., 2020; Henn et al., 2022; Manson et al., 2020; Muñoz and Grant, 2022; Song et al., 2021).

VD in humans has two origins: synthesis in the skin by the action of solar ultraviolet radiation on 7-dehydrocholesterol and the diet. The active VD metabolite is 1α,25-dihydroxyvitamin D_3_ (1,25(OH)_2_D_3_, calcitriol), which results from two consecutive hydroxylations of VD, the first in the liver and the second in the kidney or in many epithelial and immune cell types (Bikle and Christakos, 2020; Feldman et al., 2014). VD actions are mediated by 1,25(OH)_2_D_3_ binding to a member of the nuclear receptor superfamily of transcription factors, the VD receptor (VDR). Upon ligand binding, VDR regulates the expression of hundreds of target genes, many of which are involved in calcium and phosphate homeostasis, bone biology, immune response, metabolism, detoxification, and cell survival, proliferation and differentiation (Bikle and Christakos, 2020; Carlberg and Muñoz, 2022; Carlberg and Velleuer, 2022; Feldman et al., 2014; Fernández-Barral et al., 2020).

Consistent with a protective role of VD against CRC, the colonic epithelium expresses high levels of VDR. Many studies in CRC cells and associated fibroblasts and in experimental animals have shown a large series of VDR-mediated VD anti-CRC effects (Carlberg and Muñoz, 2022; Carlberg and Velleuer, 2022; Feldman et al., 2014; Fernández-Barral et al., 2020; Ferrer-Mayorga et al., 2019) In support, germline deletion of *Vdr* in mice with constitutively active Wnt/β-catenin signalling, as a main driver of CRC, results in increased intestinal tumour load (Larriba et al., 2011; Zheng et al., 2012).

Notably, certain evidence links SIRT1 and VD. First, decreased SIRT1 activity has been linked to the pathogenesis of CRC (Ren et al., 2017; Strycharz et al., 2018). However, using SIRT1 level as a marker of malignancy is controversial since both increased or decreased SIRT1 tumour expression have been described in different studies (Carafa et al., 2019; Chen et al., 2014; Firestein et al., 2008), which points to a discrepancy between SIRT1 level and activity. Second, SIRT1 deacetylates the VDR to enhance its activity in kidney and bone cells (Sabir et al., 2017) and, consequently, decreased SIRT1 activity may drive VD insensitivity. Third, promoter-bound VDR increases *SIRT1* gene expression in kidney and liver cells (Yuan et al., 2022) and thus, VD deficiency may decrease SIRT1 levels leading to reduced activity. However, whether VD alters SIRT1 activity *per se* and/or SIRT1 protein expression in CRC remains unclear.

Here we reveal that 1,25(OH)_2_D_3_ induces specific SIRT1 activity at multiple levels in human CRC cells. Notably, liganded VDR post-translationally increases SIRT1 activity through auto-deacetylation. Mechanistically, SIRT1 K610 is identified as a critical player for 1,25(OH)_2_D_3-_enhanced VDR-SIRT1 interaction, SIRT1 auto-deacetylation and activation. This occurs independently of increased SIRT1 RNA and protein expression and elevated cofactor (NAD^+^) levels. Discrepant SIRT1 level and activity may explain the controversial role of SIRT1 as tumor suppressor or tumor promoter in CRC. Our results provide a new mechanistic insight into the association of VD deficiency with CRC and suggest potential therapeutic benefit for SIRT1 activators in SIRT1-positive CRC.

## RESULTS

### 1,25(OH)2D3 increases SIRT1 level via the VDR in CRC cells

The question of whether VD governs SIRT1 activity in CRC was first approached in two well characterized CRC cell lines, HCT 116 and HT-29. Immunofluorescence analyses (Fig. 1A) revealed that SIRT1 protein level doubled in response to 1,25(OH)_2_D_3_, a result confirmed by Western blotting of nuclear extracts (Fig. 1B). The increased expression of SIRT1 protein by 1,25(OH)_2_D_3_ could be explained by a 2-fold upregulation of SIRT1 RNA revealed by RT-qPCR (Fig.1C). The induction of *CYP24A1* (Fig. 1D), a major VDR target gene in many cell types served as control for 1,25(OH)_2_D_3_ activity.

**Fig. 1.**
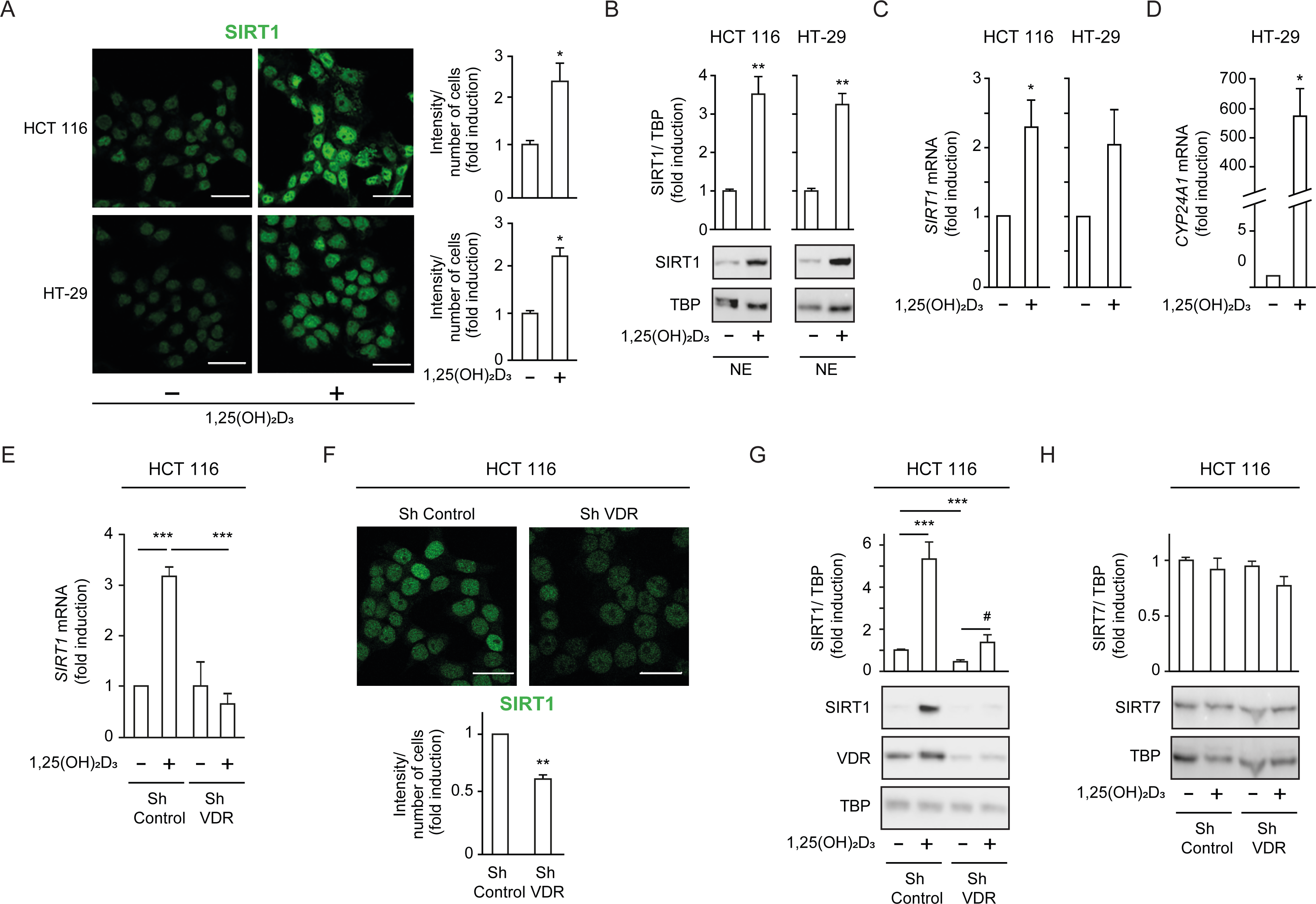
1,25(OH)_2_D_3_ increases SIRT1 levels in CRC cells and VDR is required to ensure basal levels. (A)-(D) HCT 116 or HT-29 CRC cells cultured under standard conditions and where indicated, treated with 1α,25-dihydroxyvitamin D_3_ (1,25(OH)_2_D_3_), 100 nM, added 24 hours before harvesting. (E)-(H) ShVDR cells were derived from HCT 116 by stable and specific knock-down of vitamin D receptor (VDR) using specific shRNA and are compared to ShControl cells that contain normal VDR levels (Larriba et al., 2011). Cell extracts were fractionated in all cases. (A) Confocal imaging of SIRT1 (green) in CRC (HCT 116 and HT-29) cells and fluorescence intensity quantification of 3 independent experiments using ImageJ software. For each experiment 3 different fields were evaluated per slide. Scale bars:25µm. (B) Western-blot analysis of 1,25(OH)_2_D_3_ effects on levels of SIRT1 in nuclear extracts (NE) from indicated CRC cells. Representative blots and statistical analysis using TBP as loading controls. (C)-(E) RT-qPCR analysis of the effect of 1,25 (OH)_2_D_3_ in indicated colon cancer cells. Values normalized with endogenous control (18S RNA) are referred as fold induction over cells without 1,25(OH)_2_D_3_. (C) Effect of 1,25 (OH)_2_D_3_ on SIRT1 gene expression. (D) validation of 1,25 (OH)_2_D_3_ activity on the canonical target gene cytochrome P450 family 24 subfamily A member 1, CYP24A1, in HT-29 colon cancer cells. (E) Requirement of vitamin D receptor (VDR) for 1,25 (OH)_2_D_3_ induction of SIRT1 gene expression. (F) Effect of VDR knock-down on SIRT1 protein content evaluated by confocal imaging of ShControl and ShVDR HCT 116 colon cancer cells to detect SIRT1 (green). Scale bars represent 25 μm. On the right, quantification of fluorescence intensity using ImageJ software; for each experiment, 3 different fields were evaluated per slide. (G) Western blot analysis of the effect of 1,25 (OH)_2_D_3_ on nuclear accumulation of SIRT1 in cells depleted (ShVDR) or not (ShControl) of VDR. (H) Specificity of 1,25 (OH)_2_D_3_ effects on SIRT1 by western blot analysis of the alternative nuclear sirtuin, SIRT7. Statistical analysis of 3 independent experiments in each panel was performed by Student t-test (A) to (D) and (H) or by One-Way ANOVA (E)-(F). For (G), statistical analysis of the four groups by ANOVA is represented by * and comparison of the two indicated groups by Students t test, by #. In all panels, values represent mean ± SEM of triplicates corresponding to biological replicates; * or # p< 0.05; ** p< 0.01; *** p< 0.001.

VDR expression is the main determinant of cell responsiveness to 1,25(OH)_2_D_3_ and is downregulated in a proportion of patients with advanced CRC, implying that these patients would probably not benefit from the anticancer effects of 1,25(OH)_2_D_3_ (Ferrer-Mayorga et al., 2017; Larriba and Muñoz, 2005; Pálmer et al., 2004; Peña et al., 2005). To model CRCs patients with low VDR tumour level or with VD deficiency, we used an HCT 116-derived cell line stably depleted of VDR using shRNA (hereafter named ShVDR) (Larriba et al., 2011). Induction of *SIRT1* gene expression by 1,25(OH)_2_D_3_ was confirmed by RT-qPCR analysis in ShControl but not in ShVDR cells, although basal SIRT1 levels were similar (Fig. 1E). At the protein level, confocal imaging showed reduced SIRT1 protein in ShVDR as compared to ShControl cells (Fig. 1F), suggesting that VDR contributes to maintaining basal SIRT1 protein expression. Western blot analyses on nuclear extracts confirmed that ShControl cells responded to 1,25(OH)_2_D_3_ with increased SIRT1 protein levels in contrast to the unresponsiveness of ShVDR cells (Fig. 1G). Moreover, the upregulating effect of 1,25(OH)_2_D_3_ was specific for SIRT1, since the expression of the other nuclear sirtuin, SIRT7, was unaffected (Fig. 1H). Together, these data indicated that 1,25(OH)_2_D_3_ specifically increased SIRT1 mRNA and protein levels in CRC cell lines via VDR. This is important because SIRT1 depletion and/or sirtuin activity decay might represent an important neglected mediator of VD deficiency and a target for intervention in CRC. Whether VD controls sirtuin activity independently of sirtuin level in CRC is a critical issue that deserved further exploration.

### 1,25(OH)2D3 induces SIRT1 deacetylase activity in CRC cells

Evaluation of global sirtuin activity in the nuclei of CRC cells revealed that treatment with 1,25(OH)_2_D_3_ increased NAD^+^-dependent deacetylase activity 1.5-fold as compared to control untreated cells (Fig. 2A). NAD^+^ is a cofactor required for sirtuin activity. Metabolic control of the level of NAD^+^ represents an important node for sirtuin regulation and could explain the induction of general sirtuin activity by 1,25(OH)_2_D_3_. Indeed, nuclear NAD^+^ level doubled in response to 1,25(OH)_2_D_3_ in human CRC cells (Fig. 2B). However, whether 1,25(OH)_2_D_3_ could specifically increase SIRT1 activity remained unknown.

**Fig. 2.**
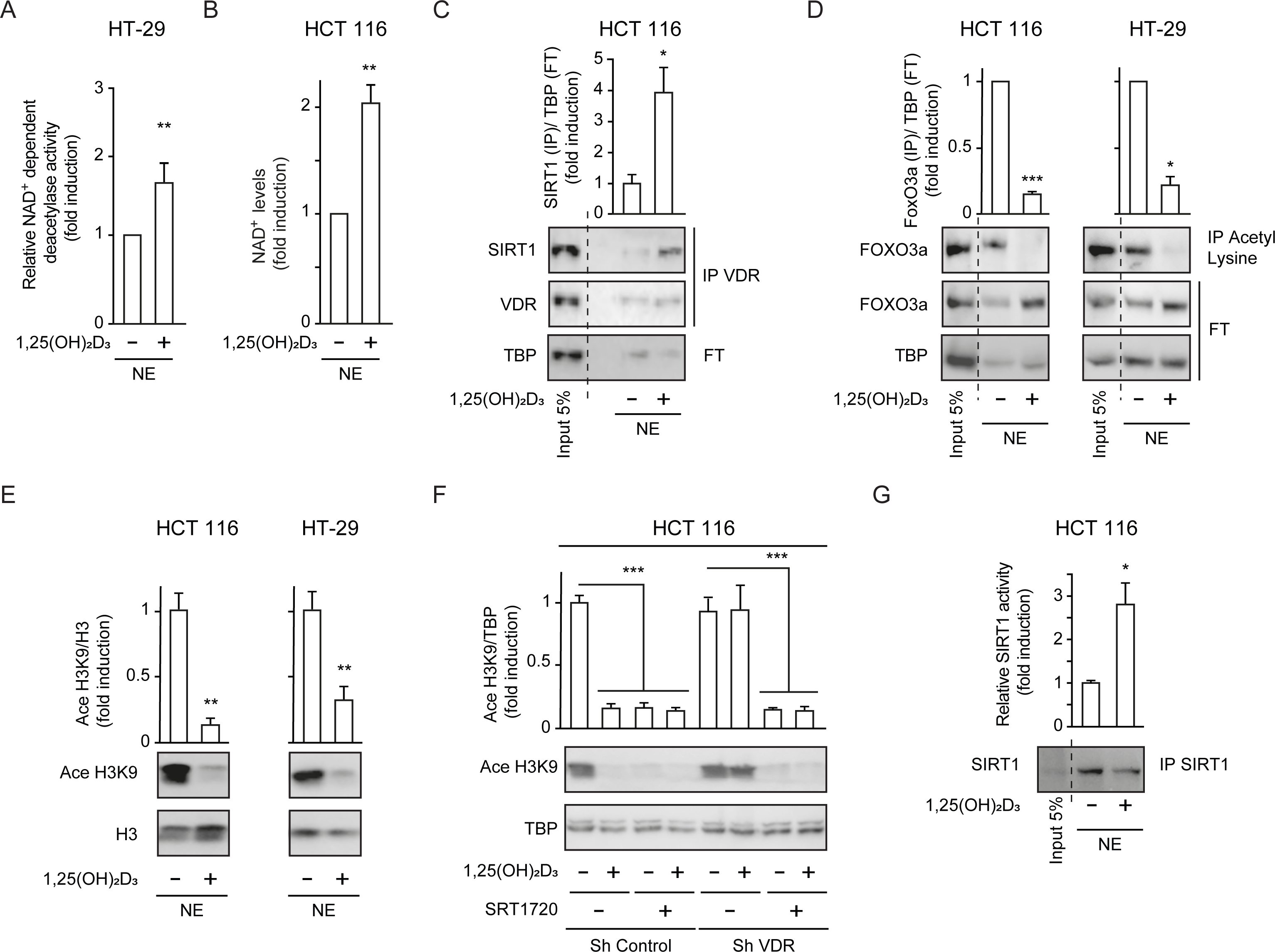
1,25(OH)_2_D_3_ induces specific SIRT1 deacetylase activity independently of levels, in CRC cells. Cells were cultured as in Fig 1. (A) and (G) NAD^+^-dependent deacetylase activity, (B) Evaluation of nuclear NAD^+^ levels. (C) to (F) Western blot analysis using TBP or H3 as loading controls. Representative western blots and statistical analysis of 3 independent experiments. (A) NAD^+^-dependent deacetylase activity measured on nuclear extracts of HT-29 cells. Relative luciferase units (RLU) were calculated as fold induction relative to the corresponding control. (B) Evaluation of nuclear NAD^+^ levels in HT-29 CRC cells in response to 1,25(OH)_2_D_3_, with data expressed as fold induction over the control, untreated cells. (C) 1,25(OH)_2_D_3_ effects on the interaction between vitamin D receptor (VDR) and SIRT1. Input (5%) and flow through (FT) are shown (D) 1,25(OH)_2_D_3_ effects on the acetylation status of the SIRT1 substrate FoxO3a in nuclear extracts immunoprecipitated with anti-acetyl lysine antibodies. Input (5%) and flow through (FT) are shown. (E) 1,25(OH)_2_D_3_ effects on the nuclear acetylation of the SIRT1 substrate H3K9 (Ace H3K9). (F) Requirement of VDR for deacetylation of Ace H3K9 in response to 1,25(OH)_2_D_3_ (G) Quantification of NAD^+^-dependent deacetylase activity on SIRT1 immunoprecipitates from nuclear extracts (NE) of HCT 116 CRC cells treated or not with 1,25(OH)_2_D_3._ Values represent mean ± SEM of n=3 independent experiments. Statistical analysis by Student t test, n ≥ 3 biological replicates and values represent mean ± SEM; *p<0.05; **p<0.01; *** p<0.001.

Liganded VDR interacts with SIRT1 in kidney and bone cells (Sabir et al., 2017) suggesting that 1,25(OH)_2_D_3_ might alter SIRT1 expression or activity via post-transcriptional mechanisms. Since protein-protein interactions regulate stability and activity of proteins, the ability of 1,25(OH)_2_D_3_ to promote VDR-SIRT1 interaction in CRC cells was evaluated by co-immunoprecipitation assays. The results revealed that 1,25(OH)_2_D_3_ potently (4-fold) enhanced VDR/SIRT1 complex formation (Fig. 2C). To test if this interaction alters SIRT1 activity, the acetylation status of two specific SIRT1 substrates (FOXO3a and H3K9) was evaluated as a read out of SIRT1 deacetylase activity in response to 1,25(OH)_2_D_3_. Acetyl-lysine immunoprecipitation followed by Western blotting demonstrated efficient deacetylation of FOXO3a in response to 1,25(OH)_2_D_3_ (Fig. 2D). Likewise, the level of acetylated histone H3K9 (Ace H3K9) dramatically decreased in response to 1,25(OH)_2_D_3_ (Fig. 2E), indicating specific activation of SIRT1 by 1,25(OH)_2_D_3_. The effect of 1,25(OH)_2_D_3_ was mediated by VDR since ShVDR cells did not show Ace H3K9 deacetylation in response to 1,25(OH)_2_D_3_, whereas ShControl cells maintained the response (Fig. 2F). Importantly, the specific SIRT1 activator SRT1720 reduced Ace H3K9 levels in both ShControl and ShVDR cells, thus mimicking 1,25(OH)_2_D_3_ action independently of VDR (Fig. 2F).

The ability of 1,25(OH)_2_D_3_ to increase protein deacetylation mediated by SIRT1 could be driven either by elevated SIRT1 level or by increased SIRT1 activity. To distinguish between these two possibilities, we performed an *in vitro* deacetylase assay using equivalent amounts of immunoprecipitated SIRT1 from cells treated or not with 1,25(OH)_2_D_3_. SIRT1-specific deacetylase activity was enhanced 3-fold in cells treated with 1,25(OH)_2_D_3_ despite similar levels of immunoprecipitated SIRT1 (Fig. 2G).

Together, these results indicated that 1,25(OH)_2_D_3_ induces nuclear SIRT1 activity. This implies that VD deficiency and/or loss of tumour VDR expression (both observed in a proportion of CRC patients) (Evans et al., 1998; Ferrer-Mayorga et al., 2019; Giardina et al., 2015; Larriba et al., 2013, 2009; Pálmer et al., 2004; Peña et al., 2005) may negatively impact SIRT1 deacetylase activity and prompted us to examine samples of CRC patients.

### CRC biopsies exhibit discrepancies between SIRT1 protein levels and deacetylase activity

Changes in SIRT1 RNA and protein expression and enzymatic activity were analysed in samples of CRC patients. First, we compared available data (TCGA-COAD database) from 160 pairs of human colon adenocarcinoma samples matched with their normal adjacent tissue. TNM plot analysis revealed that both *VDR* and *SIRT1* gene expression decreased from normal to tumour colonic tissue in a highly significant fashion (*P*<0.001) (Fig. 3A-B).

**Fig. 3.**
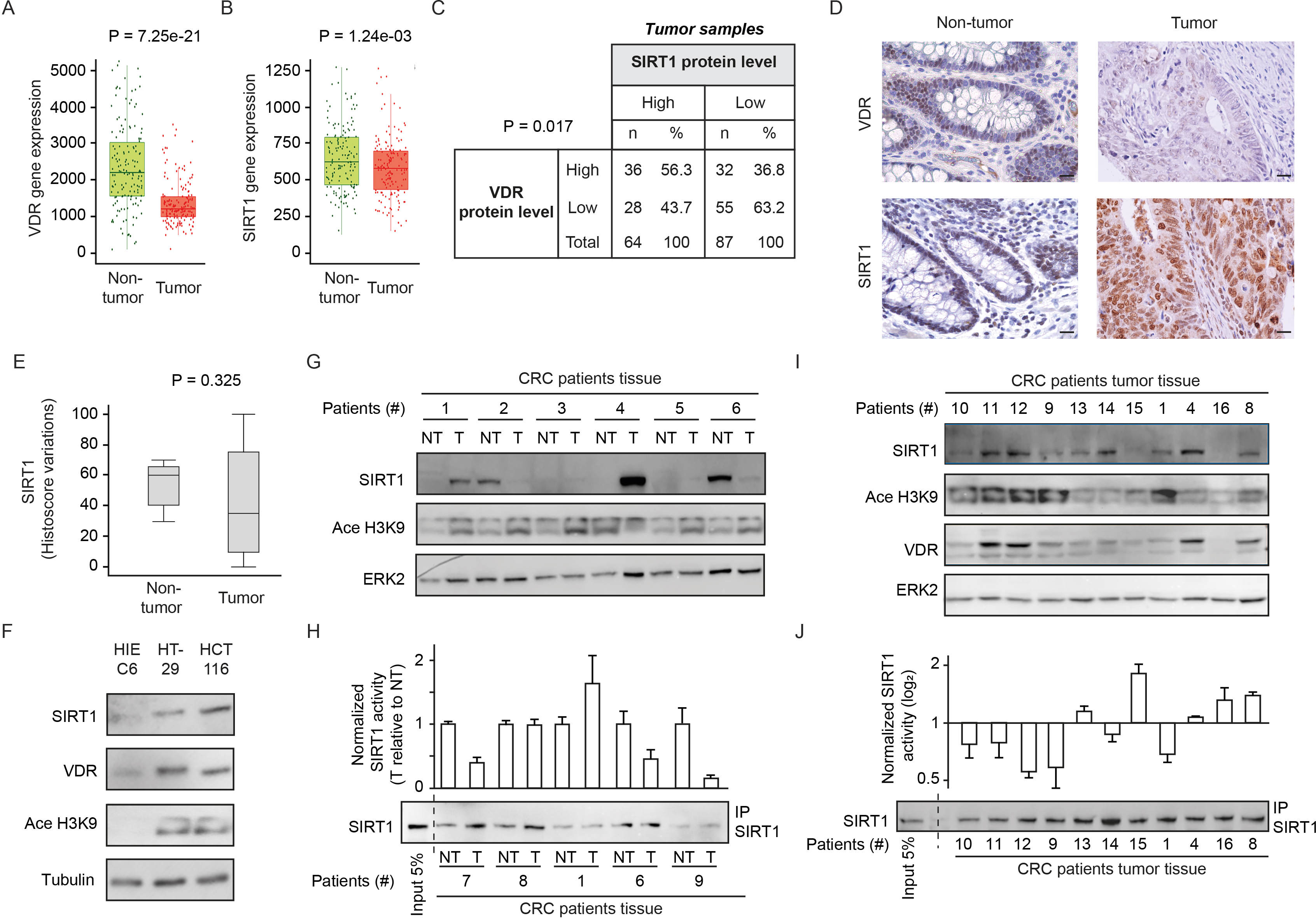
CRC biopsies exhibit discrepancies between SIRT1 protein levels and deacetylase activity. (A)-(B) TNM plot analysis of VDR (A) or SIRT1 (B) gene expression in 160 paired samples of non-cancer adjacent tissue (normal) versus adenocarcinoma tissue (tumour). Data was collected from TCGA-COAD database. SIRT1 or VDR expression is shown as transcripts per million (TPM) in log2. Tumour and control samples were compared with Mann-Whitney U test. The statistical significance cut off was set at p < 0.01. (C) Association analysis between VDR and SIRT1 protein levels in human colon cancer samples, stage 2. Cut-off points to separate proteins between high-or low-expression levels were for SIRT1 expression median of Histoscore, and negative *versus* positive staining for VDR expression. Statistical analysis was performed with Chi-square test. (D) Representative micrographs at 40X from immunostaining for vitamin D receptor (VDR) or SIRT1 on human samples of healthy colon or CRC patients. Scale bars: 20µm. Immunostainings were revealed with DAB (diaminebenzidine) and thus, positiveness is highlighted by light or dark brown according to their low or high protein expression. Counterstaining with hematoxilin stains nuclei in dark blue and cytoplasm in light blue. (E) Profile for SIRT1 content from healthy to colon cancer human samples. The profile was obtained from Histoscore variations. Tumour and control samples were compared with Mann-Whitney U test. The statistical significance cut off was set at p < 0.01. (F) Western-blot analysis of the levels of SIRT1 and its substrate Ace H3K9 in total lysates from intestinal healthy (HIEC6) or CRC cells. Representative western blot with ERK as loading control. (G) Western-blot analysis from human patient samples, with patient number indicated at the top. CRC samples (T) and adjacent nontumor tissue (NT) were probed for SIRT1 and its substrate Ace H3K9 (as in F). Frozen samples were obtained from HUFA patients. (H) NAD^+^-dependent deacetylase activity on immunoprecipitated SIRT1 from patient lysates. Values for T and NT samples were corrected according to immunoprecipitated SIRT1 levels shown on the western underneath the graph. Values for T samples were referred to values of NT and data (duplicates) are expressed as fold induction. Patient number is indicated at the bottom. (I) Western-blot analysis for SIRT1 and its substrate Ace H3K9 levels in total lysates from alternative human CRC samples (as in G). Patient number is indicated at the top. (J) NAD^+^-dependent deacetylase activity on immunoprecipitated SIRT1 from patient lysates presented in (I). Levels of immunoprecipitated SIRT1 are presented in the bottom panel. SIRT1 deacetylase activity in patients is presented as fold induction, referred to the first patient sample (patient #10). Patient number is indicated at the bottom.

Immunohistochemical analysis of human colon tumor tissue microarrays (TMAs) from the Biobank of Hospital Clínico San Carlos (HCSC, Madrid) revealed a significant (*P*=0.017) positive association between the levels of VDR and SIRT1 proteins (Fig. 3C). An additional TMA containing healthy colon samples was used to compare tumor and nontumor tissues. The level of VDR protein was usually lower in tumour samples (Fig. 3D), while SIRT1 protein expression showed high variability between tumors and high cellular intratumor heterogeneity (Fig. 3D-E). Globally, TMA data suggested that SIRT1 protein levels in tumor samples (Fig. 3E), are subjected to high variability, which hampers statistical significance even though for SIRT1 gene expression, differences were statistically significant as shown in Figure 3B.

Global acetylation also showed high cellular intratumor heterogeneity and variability between samples without statistically significant differences between tumor and nontumor samples (data not shown), showing that is not a valid, specific read out of SIRT1 activity. As an alternative approach, the levels of SIRT1 protein and its substrate Ace H3K9 were compared first in intestinal cell lines and then in human tumours. Surprisingly, SIRT1 protein level increased in CRC cells as compared to normal HIEC6 cells (Fig. 3F). However, increased SIRT1 levels coincided with elevated Ace H3K9 (Fig. 3F), indicating SIRT1 inactivation in CRC cells. The results in cell lines encouraged us to evaluate the potential discordance of SIRT1 and Ace H3K9 levels in fresh-frozen samples from paired tumor and nontumor colonic tissue of patients from Hospital Universitario Fundación Alcorcón (HUFA). All tissues were evaluated to have at least 85% of tumor cells by immunohistochemistry (IHC). Western blotting allowed simultaneous detection of SIRT1 and Ace H3K9. Notably, whereas the level of SIRT1 protein variably increased or decreased from nontumor to tumor tissues in different patients, acetylation of its specific substrate H3K9 consistently increased in tumor samples (Fig. 3G), reflecting SIRT1 inactivation as found in carcinoma cell lines (Fig. 3F). *In vitro* deacetylation assays performed on SIRT1 immunoprecipitated from SIRT1-positive patient samples confirmed a common decrease of SIRT1 activity in colon tumor tissue (Fig. 3H). Western blotting illustrated the variability of SIRT1 level among tumors and its correlation with VDR level (Fig. 3I), while Ace H3K9 levels are generally high, but variable. Importantly, in vitro deacetylation assays on SIRT1 immunoprecipitated from the same samples (Fig. 3J) confirmed Ace H3K9 as a *bona fide* read out of SIRT1 inactivation. The discrepancy between the levels of SIRT1 protein expression and activity in tumor samples may solve the controversy previously reported regarding the use of SIRT1 as a tumor biomarker and poses the question of how 1,25(OH)_2_D_3_ specifically enhances SIRT1 deacetylase activity.

### 1,25(OH)2D3 induction of SIRT1 activity is mediated through auto deacetylation

Since SIRT1 catalyses its auto deacetylation as a mechanism to increase its deacetylase activity towards other substrates (Fang et al., 2017) and 1,25(OH)_2_D_3_ facilitates VDR-SIRT1 interaction, we explored the possibility that liganded VDR increases SIRT1 activity by facilitating SIRT1 auto deacetylation. To evaluate SIRT1 acetylation status upon exposure to 1,25(OH)_2_D_3_, nuclear extracts from CRC cells challenged or not with 1,25(OH)_2_D_3_ were immunoprecipitated using anti-acetyl-lysine antibodies and subjected to Western blotting. 1,25(OH)_2_D_3_ greatly reduced the nuclear pool of acetylated SIRT1 (Fig. 4A). Remarkably, the specific SIRT1 activator SRT1720 also drove SIRT1 deacetylation in HCT 116 CRC cells (Fig. 4B). Consistently, ShVDR cells did not reduce acetylated SIRT1 in response to 1,25(OH)_2_D_3_ but retain their response to SRT1720 (Fig 4C). These results indicated that 1,25(OH)_2_D_3_ activation of SIRT1 is mediated through SIRT1 deacetylation.

**Fig. 4.**
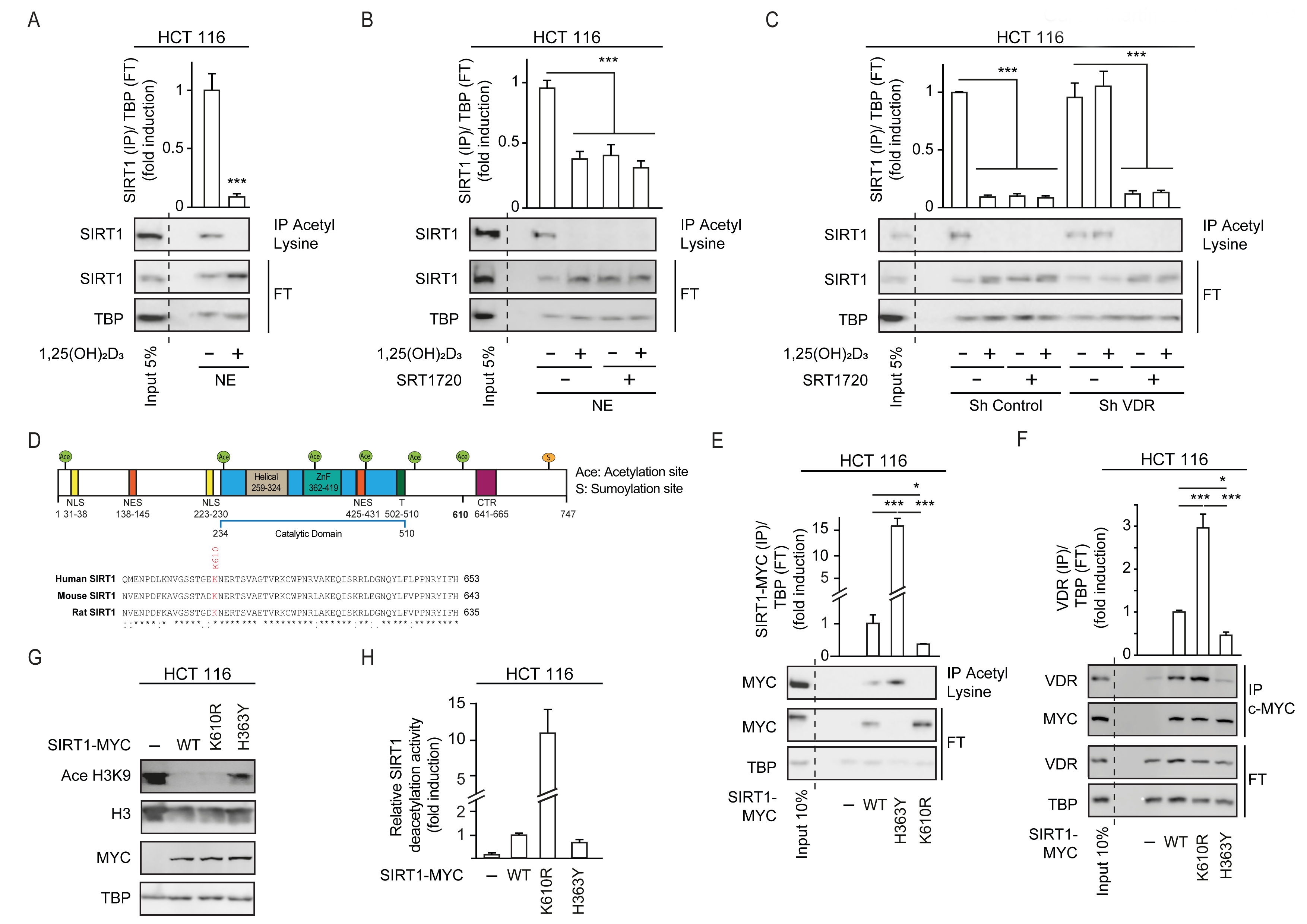
1,25(OH)_2_D_3_ induces SIRT1 activity through auto deacetylation. HCT 116 CRC cells were cultured under standard conditions in DMEM containing LiCl (40 mM) to mimic Wnt signalling before addition of 1,25(OH)_2_D_3_, 100 nM or SRT1720 (20 μM) for the last 24 h; after fractionation nuclear extracts (NE) were used. (A)-(C) and (E)-(F) Immunoprecipitation (IP) using anti-acetyl lysine antibody from nuclear extracts (NE) of indicated cells and western-blot analysis to evaluate changes in acetylation of SIRT1; TBP was used in the Flow Through (FT) as loading control. Representative western-blots and statistical analysis are shown in all panels. (A) Effect of 1,25(OH)_2_D_3_ on acetylation of SIRT1. (B) Effect of SRT1720 on acetylation of SIRT1. (C) Requirement of VDR for 1,25(OH)2D_3_ or SRT1720 effects on acetylation of SIRT1 in Sh Control and ShVDR HCT 116 cells. (D) SIRT1 scheme with functional domains and putative acetylation sites (top) and sequence alignment. Conserved putative acetylation targets for activation/inactivation of SIRT1 shown at the bottom. (E) Acetylation status of exogenous SIRT1 wild type (WT) or mutants (H363Y or K610R). HCT 116 were transiently transfected with pcDNA expression vectors: empty (-), myc-tagged SIRT1 wild type (WT) or mutants: H363Y inactive or K610R, 48 h before harvesting. Immunoprecipitation using anti-acetyl-lysine antibodies and western blot detection using myc antibodies. Representative western-blot and statistical analysis; MYC and TBP in the flow through (FT) serve as loading controls. (F) Western-blot analysis of the interaction of SIRT1 wild type and mutants, with vitamin D receptor (VDR). Expression and analysis as in (E). (G) Western-blot analysis of the effect of SIRT1 mutant expression on Ace H3K9. (H) *In vitro* NAD^+^-dependent deacetylase activity assays on equivalent amounts of exogenous SIRT1 wild type (WT), K610R and inactive H363Y mutants immunoprecipitated using anti-myc antibodies. Statistical analysis in all panels except (D) and (G) by One Way ANOVA of n>3 independent experiments. Values represent mean ± SEM; *p<0.05; **p<0.01; *** p<0.001.

Next, we sought to get further insight into the mechanism of regulation of SIRT1 activity by 1,25(OH)_2_D_3_. As the acetyltransferase EP300 acetylates and inactivates SIRT2 (Han et al., 2008) and is critical for CRC cell signalling (Chocarro-Calvo et al., 2013; Gutiérrez-Salmerón et al., 2020), we used the ASEB engine (Wang et al., 2012) to search for SIRT1 residues potentially acetylated by EP300. This approach identified K610, a lysine conserved from mouse to human, as a potential EP300 target (Fig. 4D). The role of K610 in the regulation of SIRT1 activity was assessed by generating a non-acetylable K610R mutant, which should be resistant to inactivation through acetylation. We compared the enzymatic activity of K610R SIRT1 with that of wild type (WT) and inactive H363Y SIRT1 (Vaziri et al., 2001). Nuclear extracts from cells transfected with MYC-tagged SIRT1 WT or mutants were immunoprecipitated using anti-acetyl-lysine antibodies. Western blotting using an anti-MYC antibody revealed that exogenous WT SIRT1 was acetylated (Fig. 4E), as it was endogenous SIRT1 (Fig 4A). Of note, catalytically inactive H363Y SIRT1 was highly acetylated, in striking contrast with the lack of acetylation of the K610R SIRT1 mutant (Fig. 4E), This highlights the importance of K610 for the regulation of SIRT1 activity.

Since interaction with VDR was important for SIRT1 deacetylation activity, the ability of each mutant to interact with VDR was compared in immunoprecipitation assays. The results revealed that the K610R SIRT1 mutation facilitated the interaction with VDR, whereas the inactive H363Y SIRT1 showed low interaction with VDR (Fig 4F). This suggested that acetylation of SIRT1 might interfere with or destabilize its interaction with VDR, and that the ability of 1,25(OH)_2_D_3_ to favour VDR-SIRT1 interaction may allow SIRT1 auto deacetylation, and thus enhance its activity. The activity of mutant SIRT1 enzymes was first inferred from the level of Ace H3K9, which was lower in cells expressing K610R SIRT1 (Fig 4G). *In vitro* NAD^+^-dependent deacetylase assays were next used to compare the activity of WT and mutant SIRT1. Using equivalent amounts of immunoprecipitated MYC-tagged enzymes, we found a 10-fold higher deacetylase activity for the K610R mutant than for WT SIRT1, whereas as expected the H363Y inactive mutant exhibited very low activity as expected (Fig. 4H).

### SIRT1 activation mimics the antiproliferative effects of 1,25(OH)_2_D_3_ in CRC cells unresponsive to VD

Collectively, our data indicated that the acetylation status of SIRT1 K610 is critically controlled and governs SIRT1 activity in the nuclei of CRC cells. By promoting VDR-SIRT1 interaction, 1,25(OH)_2_D_3_ facilitates SIRT1 auto deacetylation and enhances SIRT1 activity. These results suggest that SIRT1 is an effector of 1,25(OH)_2_D_3_ and prompted us to ask whether the pharmacological activation of SIRT1 would rescue CRC cells that are unresponsive to 1,25(OH)_2_D_3_.

The anticancer effects of 1,25(OH)_2_D_3_ are exerted at least partially through the inhibition of tumor cell proliferation (Carlberg and Muñoz, 2022; Muñoz and Grant, 2022). 1,25(OH)_2_D_3_ strongly reduced the percentage of CRC cells in S-G_2_M cell-cycle phases (Fig. 5A) and consequently, it lengthened the cell cycle and reduced CRC cell proliferation. We aimed to analyse the effect of the modulation of SIRT1 activity on the proliferation of CRC cells and on the antiproliferative action of 1,25(OH)2D3. Inhibition of general sirtuin activity with Nicotinamide (NAA) abolished the antiproliferative effects of 1,25(OH)_2_D_3_ on CRC cells (Fig. 5B). Moreover, activation of SIRT1 by the small molecule SRT1720 mimicked the response to 1,25(OH)_2_D_3_ and efficiently reduced the percentage of cells in S-G_2_M cell cycle phases (Fig. 5C). These data indicate that induction of SIRT1 activity mediates 1,25(OH)_2_D_3_-driven cell cycle lengthening in CRC cells.

**Fig. 5.**
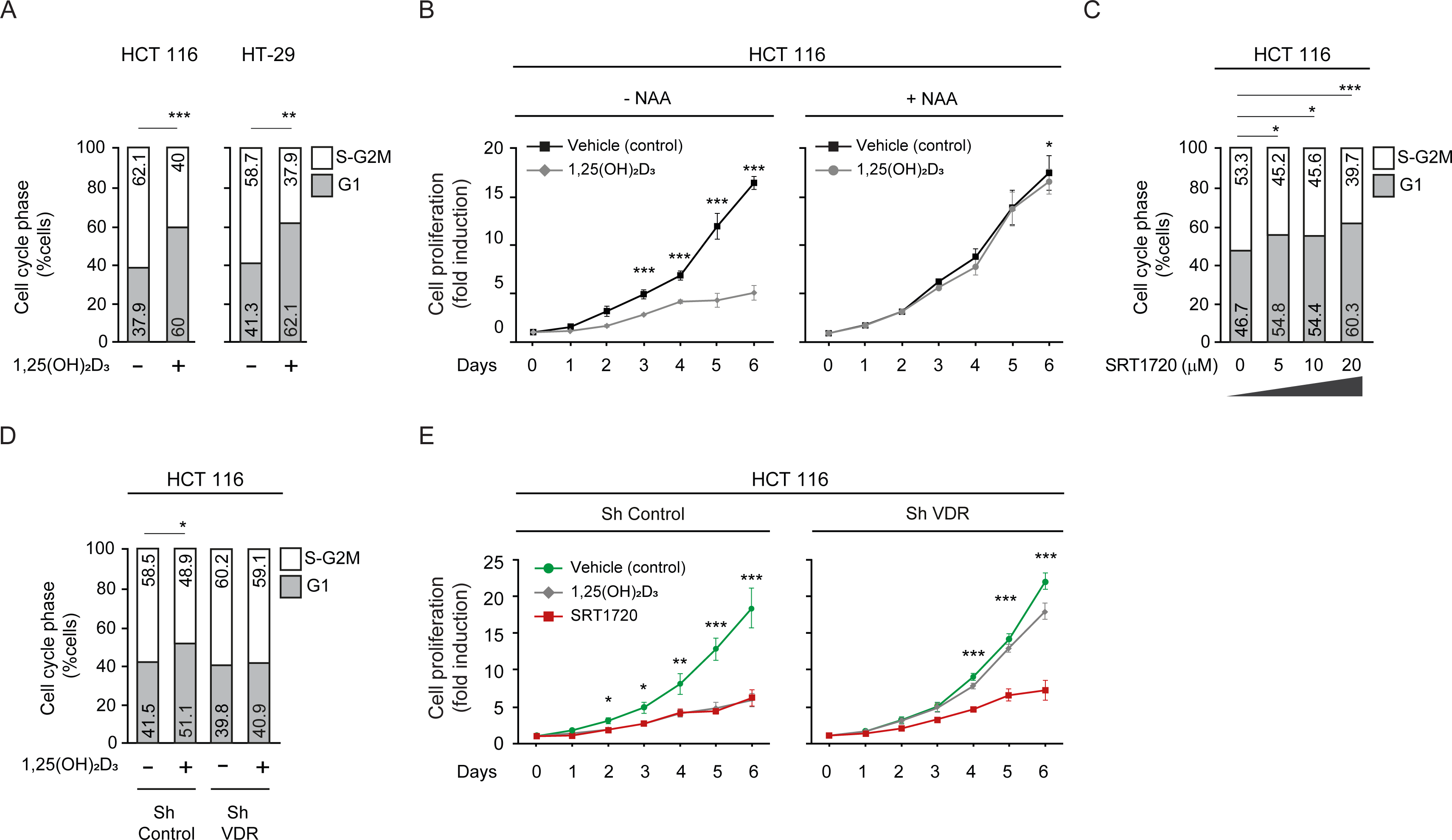
SIRT1 activation rescues antiproliferative effects of 1,25(OH)_2_D_3_ in unresponsive CRC cells. Cells were cultured as in previous figures and treatments with NAA (300 uM), SRT1720 (20μM) and 1,25 (OH)_2_D_3_ (100 nM) also were for the last 24 h. (A) Flow cytometry analysis of cell cycle from indicated cells treated (+) or not (-) with 1,25(OH)_2_D_3_. Numbers correspond to the percentage of cells in the indicated phases expressed as mean ± SEM. (B) Effect of NAA on the 1,25(OH)_2_D_3_-driven blockades of HCT 116 CRC cell proliferation. (C) Effects of SRT1720 on cell cycle of HCT 116 CRC cells by flow cytometry analysis. Numbers correspond to the percentage of cells in the indicated phases expressed as mean ± SEM. (D) Flow cytometry analysis of cell cycle response to 1,25 (OH)_2_D_3_ in ShControl and ShVDR HCT 116 cells. Numbers correspond to the percentage of cells in the indicated phases expressed as mean ± SEM. (E) Proliferation curves of ShControl or ShVDR HCT 116 colon cancer cells in response to 1,25 (OH)_2_D_3_ (100 nM) (grey), compared to control with vehicle (green) or SRT1720 (red). Statistical analysis by One-Way ANOVA of 3 independent experiments; values represent mean ± SEM of triplicates; * p< 0.05; ** p< 0.01; *** p< 0.001.

As expected, in contrast with ShControl cells, ShVDR CRC cells did not extend the cell cycle in response to 1,25(OH)_2_D_3_ (Fig. 5D). Accordingly, an early and strong reduction in the proliferation rate in response to 1,25(OH)_2_D_3_ was noted in ShControl cells but it was almost absent in ShVDR cells (Fig. 5E). Specific SIRT1 activation with SRT1720 reduced the proliferation of ShControl cells at a comparable magnitude to 1,25(OH)_2_D_3_ and, importantly, also reduced proliferation of 1,25(OH)_2_D_3_-unresponsive ShVDR cells (Fig.5E).

Collectively, these results show that SIRT1 acts downstream of 1,25(OH)_2_D_3_-activated VDR in the inhibition of CRC cell proliferation. They also suggest that activation of SIRT1 in CRC patients with vitamin D deficiency or with VD unresponsive tumors (conditions frequently observed in advanced CRC), could be a therapeutic strategy to effectively antagonize cell proliferation.

## DISCUSSION

Here we reveal that 1,25(OH)_2_D_3_ activates the epigenetic regulator SIRT1 by acting at multiple levels. Ligand-activated VDR post-translationally upregulates SIRT1 activity by facilitating its auto deacetylation. Mechanistically, SIRT1 K610 is identified as essential to control VDR-SIRT1 interaction and SIRT1 acetylation status and activity. Notably, both SRT1720 and 1,25(OH)_2_D_3_ activate SIRT1 through auto deacetylation. Modulation of SIRT1 activity by 1,25(OH)_2_D_3_ through deacetylation occurs in addition to increased mRNA and protein levels as well as to an elevation of cofactor (NAD^+^) availability. Thus, 1,25(OH)_2_D_3_ activates SIRT1 acting at multiple levels, from gene expression to activity, and these findings broaden our understanding of how VD deficiency is associated with CRC.

We also show that SIRT1 deacetylase activity mediates the anti-proliferative action of 1,25(OH)_2_D_3_ in CRC, which is in line with the recent proposal that SIRT1 activity opposes several cancer cell hallmarks (Yousafzai et al., 2021). In cases of VD deficiency and/or unresponsiveness such as advanced colorectal tumor lacking VDR expression due to transcriptional repression by SNAIL1/2 (Pálmer et al., 2004; Larriba et al., 2009), the activation of SIRT1 by compounds other than 1,25(OH)_2_D_3_ may be especially relevant. Clinical trials that explore SIRT1 activators in CRC are rare (https://clinicaltrials.gov). Some trials are designed to study doses and pharmacokinetics of resveratrol, a known sirtuin activator that has many other targets (Patel et al., 2011). Currently there are not known clinical trials using SRT1720 or alternative SIRT1 specific activators.

Consideration of SIRT1 protein level as a prognostic marker for cancer has received much attention, but conflicting results reporting both, tumor SIRT1 increases and decreases raised controversy (Ren et al., 2017; Wu et al., 2017). Accordingly, some studies propose SIRT1 either as tumour suppressor or as tumor promoter in CRC (Carafa et al., 2019; Ren et al., 2017). Importantly, SIRT1 protein level might not always reflect SIRT1 activity, and some tumors could possibly increase SIRT1 protein expression to counteract inactivation. Our results, which indicate the existence of discrepancies between SIRT1 levels and activity in human colorectal tumors, may help to resolve the controversy. Interestingly, alternative posttranslational SIRT1 modifications may have a similar outcome. For example, proteolytic cleavage of SIRT1 induced by TNFα in osteoarthritis renders accumulation of a catalytically inactive 75 kDa resistant fragment (Oppenheimer et al., 2012) that could be detected by immunohistochemistry. Since inflammation (in general) and especially TNFα is an important mediator of CRC, understanding if this fragment is generated in CRC patients and whether its presence would also lead to decreased SIRT1 activity in samples strongly positive for SIRT1 remains as an interesting unexplored issue.

The ability of 1,25(OH)_2_D_3_ to activate SIRT1 may have further major implications for physiological and pathological processes other than CRC since SIRT1 target proteins control several processes such as mitochondria physiology, apoptosis, and inflammation (Strycharz et al., 2018). Thus, 1,25(OH)_2_D_3_ may not only regulate gene expression via direct transcriptional regulation of VDR-bound targets, but it can secondarily impact a wide range of events and processes via the control of SIRT1 activity. This may explain how 1,25(OH)_2_D_3_ regulates the activity of FOXO3a, an important modulator of cell proliferation and the immune response that is targeted by SIRT1 (An et al., 2010). SIRT1 also modulates the activity of the tumour suppressor p53, whose mutation is a relatively early event in a high proportion of CRCs (Polidoro et al., 2013; Strycharz et al., 2018). In this way, SIRT1 activation greatly broaden the known vitamin D regulatory action, albeit whether 1,25(OH)_2_D_3_ directs SIRT1 activity towards specific substrates remains to be elucidated. The accumulating evidence for SIRT1 involvement in chronic inflammation (Mendes et al., 2017) suggests a potential benefit of SIRT1 activation by 1,25(OH)_2_D_3_ in chronic inflammation-linked CRC and other diseases. Moreover, the mechanism revealed here for SIRT1 activation by 1,25(OH)_2_D_3_ may be relevant for diseases associated with VD deficiency beyond CRC, including autoimmune disorders and diabetes.

In conclusion, the multilevel activation of SIRT1 deacetylase by 1,25(OH)_2_D_3_ broadens the range of known 1,25(OH)_2_D_3_ targets and positions SIRT1 as an important mediator of the protective action of VD against CRC and potentially other neoplasias and non-tumoral diseases. These data support SIRT1 activators as useful agents in situations of dysfunction of the VD system.

## MATERIALS AND METHODS

All materials and resources can be found in Table 1.

**Table 1.**
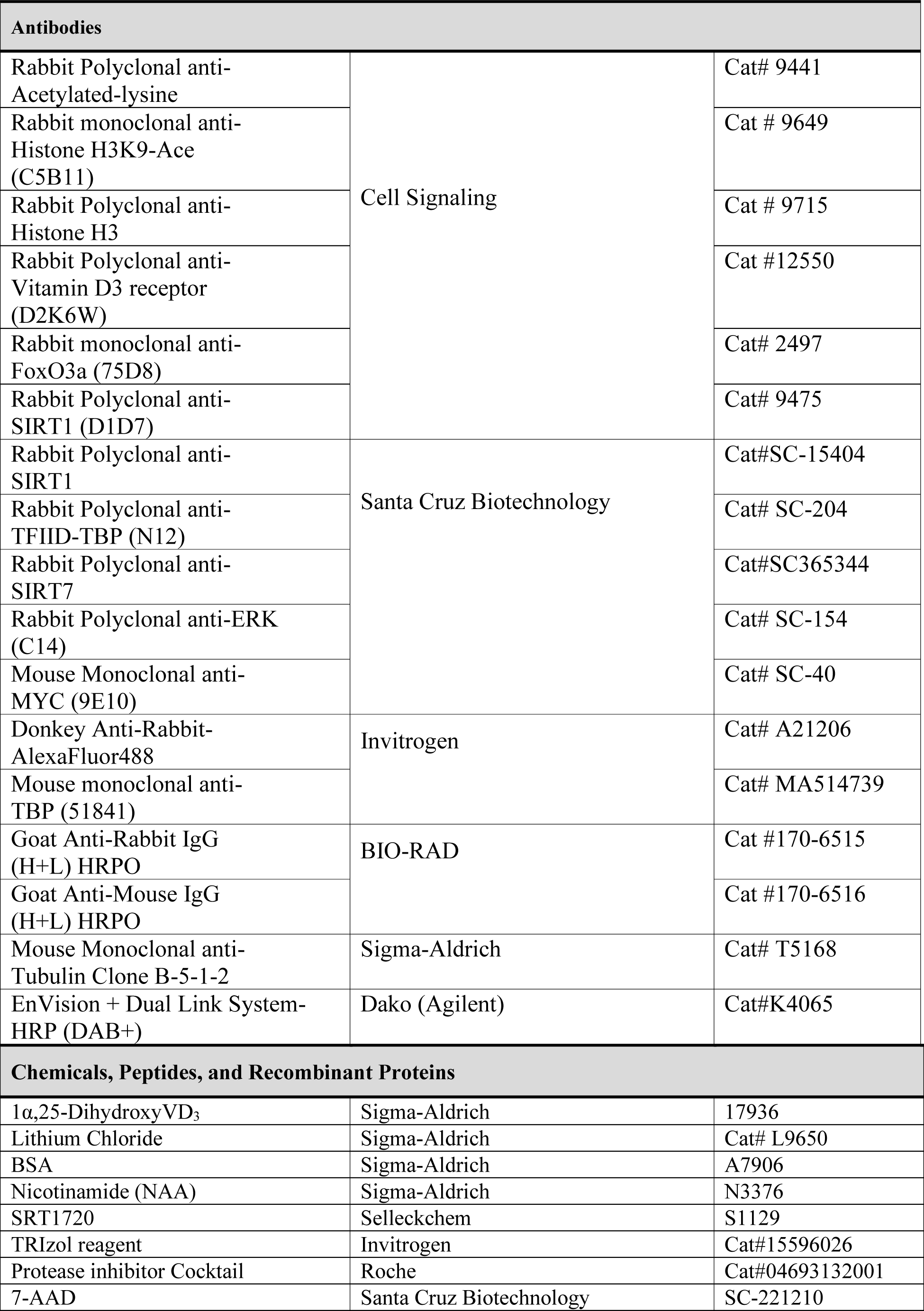
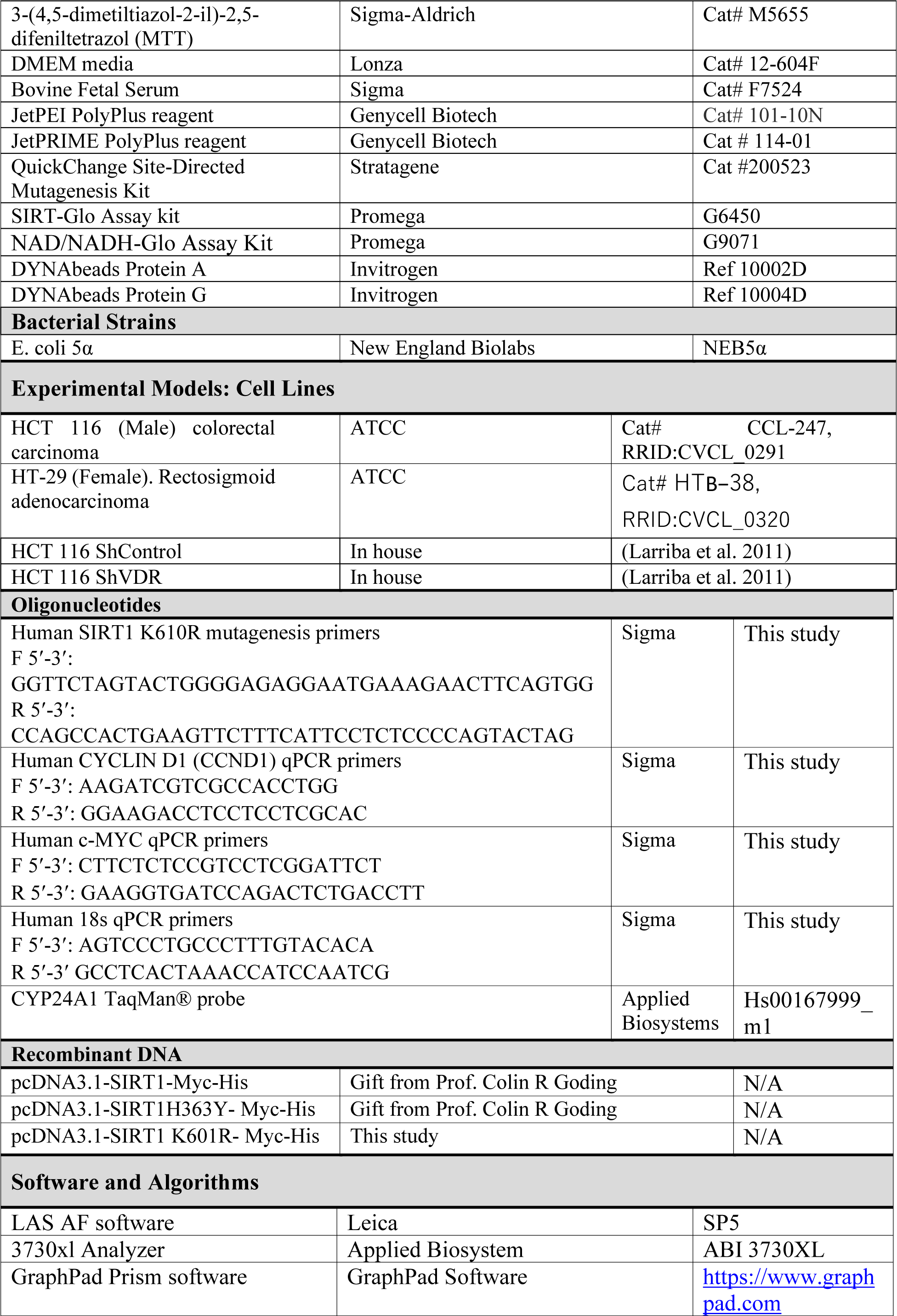
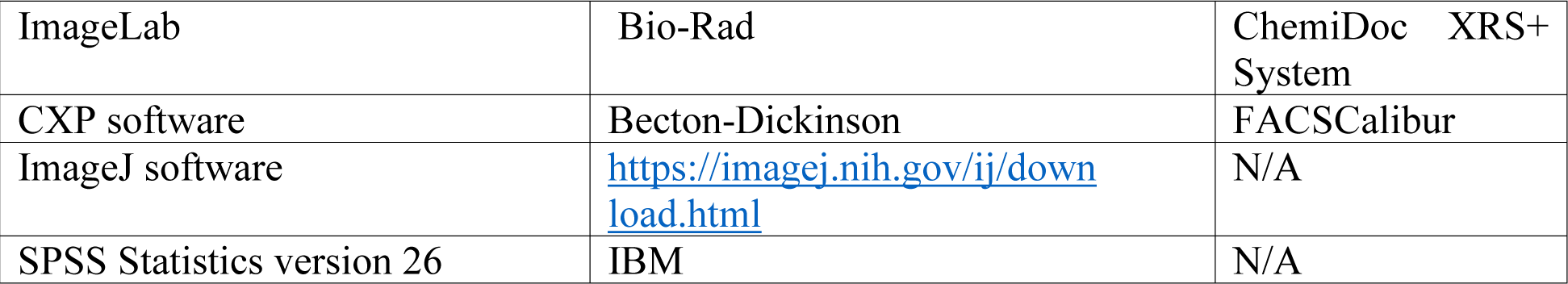
Key Materials & Resources.

### Experimental model and subject details Human samples

Patient samples were derived from surgical removal at Fundación Jimenez Diaz University Hospital, General and Digestive Tract Surgery Department: 95 patient samples diagnosed with stage II CRC. The Institutional Review Board (IRB) of the Fundación Jimenez Diaz Hospital, reviewed and approved the study, granting approval on December 9, 2014, act number 17/14. Samples from seven healthy individuals were obtained from Hospital Clínico San Carlos (HCSC). IRB-HCSC act number 21/498-E granting approval on June 25^th^, 2021. Paired fresh-frozen samples from tumor and nontumor adjacent tissue were collected from 18 patients at Hospital Universitario Fundación Alcorcón (HUFA). IRB-HUFA, act number 17/68 granting approval on June 8^th^, 2017. Colon cancer tissues were collected using protocols approved by the corresponding Ethics Committees and following the legislation. All patients gave written informed consent for the use of their biological samples for research purposes. Fundamental ethical principles and rights promoted by Spain (LOPD 15/1999) and the European Union EU (2000/C364/01) were followed. All patients’ data were processed according to the Declaration of Helsinki (last revision 2013) and Spanish National Biomedical Research Law (14/2007, July 3).

### Colorectal cell panels

Human colorectal adenocarcinoma HT-29 and HCT 116 and HCT 116 derived ShControl and ShVDR cells were cultured under standard conditions, in 5%CO2 at 37°C with DMEM containing 25 mM glucose (unless specifically indicated) and 40 mM LiCl to mimic Wnt signalling and supplemented with 10% foetal bovine serum (FBS) and 1% penicillin-streptomycin. Cells were treated as indicated for 24h. HCT 116 ShControl and ShVDR cell lines were derived previously (Larriba et al., 2011)

### Transient transfections

Cells were seeded in plates at 50% confluence for plasmid transfection, using JetPei PolyPlus reagent (Genycell Biotech), following the manufacturer’s instructions. After 24 h cells were treated as indicated for 24 h.

For Sirt1 siRNA, cells plated in six well plates at 50% confluence were transfected with JetPRIME PolyPlus reagent (Genycell Biotech) following the manufacturer’s instructions. After 2 days cells were treated 24 h with LiCl and then another 24 h with 1,25(OH)_2_D_3_ before harvesting to analyse by western blot.

### Site directed mutagenesis

Directed mutagenesis was performed using the QuickChange Site-Directed Mutagenesis Kit (Stratagene). Oligonucleotides containing the mutation (Table 1) were designed following the manufacturer’s recommendations. 2.5 U of PfuTurbo DNA polymerase was added to each sample and they were amplified by performing the following cycles: 1 denaturation cycle at 95 ° C for 1 minute, 18 amplification cycles of 30 seconds at 95 ° C, 50 seconds at 60 ° C, 1 minute / kilobase of plasmid at 68 °C and the last extension cycle 7 minutes at 68ªC. Once amplified, 10 U of Dnp I enzyme were added and incubated at 37 ° C for one hour to digest the supercoiled double-stranded DNA and eliminate the parental molecules that are not mutated.

The correct introduction of the mutation was evaluated by plasmid sequencing using the BigDye Cycle Sequencing Kit (Applied Biosystems).

### Preparation of cell extracts

#### Whole cell extracts

Cells were washed with iced PBS before extract preparation and scraped in RIPA buffer (10 mM Tris HCl [pH 7.4], 5 mM EDTA, 5 mM EGTA, 1% Tryton X100, 10 mM Na_4_P_2_O_7_ [pH 7.4], 10 mM NaF, 130 mM NaCl, 0.1% SDS, 0,5% Na-deoxycholate). After 5 min on ice, cells were pelleted (12000 rpm for 5 min, 4°C) and the supernatant was directly used as whole cell extract or frozen at −80 °C.

#### Fractionated cell extracts

After washing as before, cells were scraped in hypotonic buffer (20 mM Hepes, [pH 8.0], 10 mM KCl, 0,15 mM EDTA, 0,15 mM EGTA, 0,05% NP40 and protease inhibitors) and incubated on ice for 10 min before adding 1:2 vol of sucrose buffer (50 mM Hepes [pH=8.0], 0.25 mM EDTA, 10 mM KCl, 70% sucrose). Lysates were fractionated (5000 rpm for 5 min at 4°C) to obtain the cytoplasmic fraction in the supernatant. Nuclear pellets were washed twice with washing buffer (20 mM Hepes [pH 8.0], 50 mM NaCl, MgCl_2_ 1.5 mM, 0,25 mM EDTA, 0,15 mM EGTA, 25% glycerol and protease inhibitors), pelleted at 5000 rpm, 5 min at 4°C and resuspended in nuclear extraction buffer (20 mM Hepes[pH 8.0], 450 mM NaCl, MgCl_2_ 1.5 mM, 0,25 mM EDTA, 0,15 mM EGTA, 0,05% NP40, 25% glycerol and protease inhibitors) before centrifugation at 12,000 rpm for 5 min at 4°C to pellet and discard cell debris. The supernatants were used as nuclear fractions.

### SIRT1 Activity Assay

NAD^+^-dependent deacetylase activity was measured on nuclear extracts of HT-29 cells treated as indicated, using SIRT-Glo Assay kit (Promega), following the manufacturer’s instructions. For each reaction, 1 μg of protein was incubated with SIRT-Glo Reagent Mix at room temperature for 3 min, and the luminescence was measured using a GloMax Microplate Reader (Promega). Relative luciferase units (RLU) were calculated as fold induction relative to the corresponding control.

### Nuclear NAD_+_ abundance

The NAD/NADH-Glo Assay Kit (Promega) was used to measure NAD^+^ abundance in HCT 116 nuclear extracts. 20 μg of protein were diluted in 50 μl of lysis buffer (described for cell extracts) for each reaction and 25 μl of 0.4 N HCl were added before incubation at 60 °C for 15 minutes. This exploits the high stability of NAD^+^ under acidic conditions, while NADH breaks down. Samples were cooled down to room temperature for 10 min and neutralized with 25 µL 0.5 M Trizma®. NAD^+^ levels were then measured using a GloMax microplate reader (Promega) and following the manufacturer’s instructions.

### Immunoprecipitation

For immunoprecipitation from fractionated extracts, the hypotonic buffer was modified by adding 100mM NaCl and 0.1% NP40. For immunocomplex formation, protein A/G-coated magnetic beads (Invitrogen) were washed 3 times with the extraction buffer before coating with the primary antibody for 2 hr at 4 °C in a rotating wheel, followed by 2 washes with the same buffer to eliminates unbound antibody and then extracts were added O/N at 4°C in the rotating wheel. Immunocomplexes were washed twice and used for western blotting.

### Western blotting

Proteins from lysed cells or immunoprecipitates were denatured and loaded on sodium dodecyl sulfate polyacrylamide gels and then transferred to polyvinylidene difluoride membranes (Bio-Rad). After blocking with 5% (w/v) BSA or milk the membrane was incubated with the corresponding primary and secondary antibodies (Bio-Rad). The specific bands were analyzed using Thyphoon or ChemiDoc Imaging Systems (Bio-Rad).

### Immunoflorescence

Cells in cover slips were washed three times and fixed with 4% paraformaldehyde in PBS [pH 7.4] for 10min, washed again, permeabilized (PBS [pH7.4], 0.5% Triton X-100, 0.2% BSA) for 5min, blocked (PBS [pH7,4, 0,05% Triton X-100, 5% BSA) for 1h at room temperature and incubated with primary antibody over night at 4°C. After three washes (5 min) cells were incubated with secondary antibody for 1h at room temperature. Slides were mounted and images were acquired using a SP5 confocal microscope (Leica) with a 63x objective. Fluorescence intensity was quantified using Image J software. For each experiment, 3 different fields were evaluated per slide.

### Cell-growth curves and cell cycle

Cell growth was determined by colorimetry. For that, cells were seeded at a density of 20,000 cells per well in a Corning 12-well plate and treated as indicated for 1 to 6 days. After that cells were treated with 3-(4,5-dimethylthiazol-2-yl)-2,5-diphenyltetrazolium bromide (MTT) (Sigma-.Aldrich) in a 1:10 ratio in a culture medium and incubated at 37 °C for 3 h. In living cells, MTT is reduced to formazan, which has a purple color. The medium was removed, and the formazan was resuspended in dimethylsulfoxide (DMSO), transferred to p96 plates and analyzed by a Spectra FLUOR (Tecan) at 542 nm. Viability was measured in four independent experiments and then duplicated.

For cell cycle analysis, cells were harvested by trypsinization, washed with PBS, fixed in 70% ethanol, stained with 7-AAD (Santa Cruz Biotechnology) for 10 min at 37°C, and analyzed by flow cytometry (FACSCalibur, Becton-Dickinson). The percentages of cells at each cell cycle phase were analyzed using CXP software (Becton-Dickinson).

### RT-qPCR

Total RNA was extracted from 3 replicates of colorectal cells using TRIzol reagent (Invitrogen). Reverse transcription of 1μg of RNA was performed according to the manufacturer’s instructions Reagents and detection systems were from Applied biosystems. 18S ribosomal RNA primers served as a nonregulated control. Relative expression was calculated using the Ct method, expressed as 2^−ΔΔCt^ (Livak and Schmittgen, 2001). The PCR efficiency was approximately 100%.

### TMA, immunohistochemistry, and quantification

Tissue microarrays (TMA) containing 92 cores of stage 2 human CRC samples, were constructed using the MTA-1 tissue arrayer (Beecher Instruments, Sun Prairie) for immunohistochemistry analysis. Each core (diameter 0.6 mm) was punched from pre-selected tumour regions in paraffin-embedded tissues. We chose central areas from the tumour, avoiding foci of necrosis. Staining was conducted in 2-μm sections. Slides were deparaffinized by incubation at 60°C for 10 min and incubated with PT-Link (Dako, Agilent) for 20 min at 95°C in low pH to detect VDR, SIRT1 and acetyl lysines. To block endogenous peroxidase, holders were incubated with peroxidase blocking reagent (Dako, Agilent) and then with (1:50) dilutions of antibodies: anti-VDR, anti-SIRT1 and (1:300) anti-acetyl lysine, overnight at 4°C. All previously described antibodies presented high specificity. After that, slides were incubated for 20 min with the appropriate anti-Ig horseradish peroxidase-conjugated polymer (EnVision, Dako, Agilent). Sections were then visualized with 3,3’-diaminobenzidine (Dako, Agilent) as a chromogen for 5 min and counterstained with Harrys’ Haematoxylin (Sigma Aldrich, Merck). Photographs were taken with a stereo microscope (Leica DMi1). According to the human protein atlas (available at http://www.proteinatlas.org), a human colon tissue was used as a positive control for immunohistochemical staining to determine anti-VDR concentration and a human testis tissue for anti-SIRT1. Immunoreactivity was quantified blind with a Histoscore (H score) that considers both the intensity and percentage of cells stained for each intensity (low, medium, or high) following this algorithm (range 0–300): H score = (low%) × 1 + (medium%) × 2 + (high %) × 3. Quantification for each patient biopsy was calculated blindly by 2 investigators (MJFA and JMU). VDR and SIRT1 showed nuclear staining, whereas acetyl lysines were nuclear and extranuclear. Clinicopathological characteristics of patients are summarized in Table 2.

**Table 2.** Clinicopathological characteristics of CRC patients included in the study.

| Patients Characteristic | N (%) |  |
| --- | --- | --- |
|  | Stage II | Stage IV |
| <u>Gender</u> |  |  |
| Male | 57 (60%) | 27 (48%) |
| Female | 38 (40%) | 23 (41%) |
| N/A | 0 | 6 (11%) |
| <u>Tumor Location</u> |  |  |
| Cecum | 13 (14%) | 3 (5%) |
| Right | 25 (26%) | 10 (18%) |
| Transverse | 7 (8%) | 3 (5%) |
| Left | 5 (5%) | 2 (4%) |
| Sigma | 24 (25%) | 21 (38%) |
| Rectum | 21 (21%) | 12 (21%) |
| N/A | 0 | 5 (9%) |
| <u>Grade Primary Tumor</u> |  |  |
| Well differentiated | 18 (19%) | 15 (27%) |
| Moderately differentiated | 69 (73%) | 27 (48%) |
| Poorly differentiated | 8 (8%) | 5 (9%) |
| N/A | 0 | 9 (16%) |
| <u>Paired Liver Metastasis</u> |  | 54 (96%) |
| N/A |  | 2 (4%) |
| <u>Grade Metastasis Tumor</u> |  |  |
| Well differentiated |  | 12 (21%) |
| Moderately differentiated |  | 29 (52%) |
| Poorly differentiated |  | 4 (7%) |
| N/A |  | 11 (20%) |
| <u>Metastasis at diagnosis</u> |  |  |
| Synchronous |  | 35 (62%) |
| Metachronous |  | 21 (38%) |
| <u>Pt</u> |  |  |
| T1 | 3 (3%) | 2 (4%) |
| T2 | 30 (32%) | 5 (9%) |
| T3 | 61 (64%) | 42 (75%) |
| T4 | 1 (1%) | 4 (7%) |
| N/A |  | 3 (5%) |
| <u>pN</u> |  |  |
| N0 | 95 (100%) | 27 (48%) |
| N1 |  | 16 (29%) |
| N2 |  | 9 (16%) |
| N3 |  | 1 (2%) |
| N/A |  | 3 (5%) |

### Bioinformatics analyses

Gene expression (RNA-seq) data from the TCGA colon adenocarcinoma cohort (COAD) was downloaded from the cBioportal for Cancer Genomics (http://www.cbioportal.org) using CGDS-R (Cerami et al., 2012; Gao et al., 2013) following TCGA guidelines (http://cancergenome.nih.gov/publications/publicationguidelines). SIRT1 and VDR gene expression from samples of colorectal cancer and healthy tissue adjacent to the tumour were analysed with TNMplot.com (https://tnmplot.com/analysis/) tool (Bartha and Győrffy, 2021).

An asymptotic Spearman correlation test using original log2 expression values was used to determine the significance of the Spearman rank correlation.

### Statistical analysis

Results are presented as fold induction, mean ± SEM from three biological replicates. To determine whether calculated Hscores for protein levels were well-modelled by a normal distribution, Kolmogorov-Smirnov test was used. Association between SIRT1 and VDR protein levels was performed with Chi-square test. Comparisons between two independent groups (healthy versus tumors) were performed with Mann-Whitney U-test. For statistical association, Hscore of antigens were categorized as high-or low-expression levels using the median as a cut-off point. Tests for significance between two sample groups were performed with Student’s t test and for multiple comparisons, ANOVA with Bonferroni’s post-test.

## ACKNOWLEDGMENTS

Funding for this work was provided by the Agencia Estatal de Investigación (PID2019-104867RB-I00 / AEI/10.13039/501100011033, RTI2018-099343-B-100 and PID2021-127645OA-I00); Instituto de Salud Carlos III (CIBERONC, CB16/12/00273 and CB16/12/00326); Comunidad de Madrid (Ayudas Atracción de Talento 2017-T1/BMD-5334 and 2021-5A/BMD-20951; A385-DROPLET Young Reserchers R&D Project 2019 CAM-URJC; PRECICOLON-CM, P2022/BMD7212); Universidad Rey Juan Carlos (ADIPOMELM, Proyecto Puente de Investigación 2020).

We thank Lucille Banham for her valuable assistance in the preparation of the English manuscript and María Gutiérrez Salmerón for help making figures.

## Additional Information

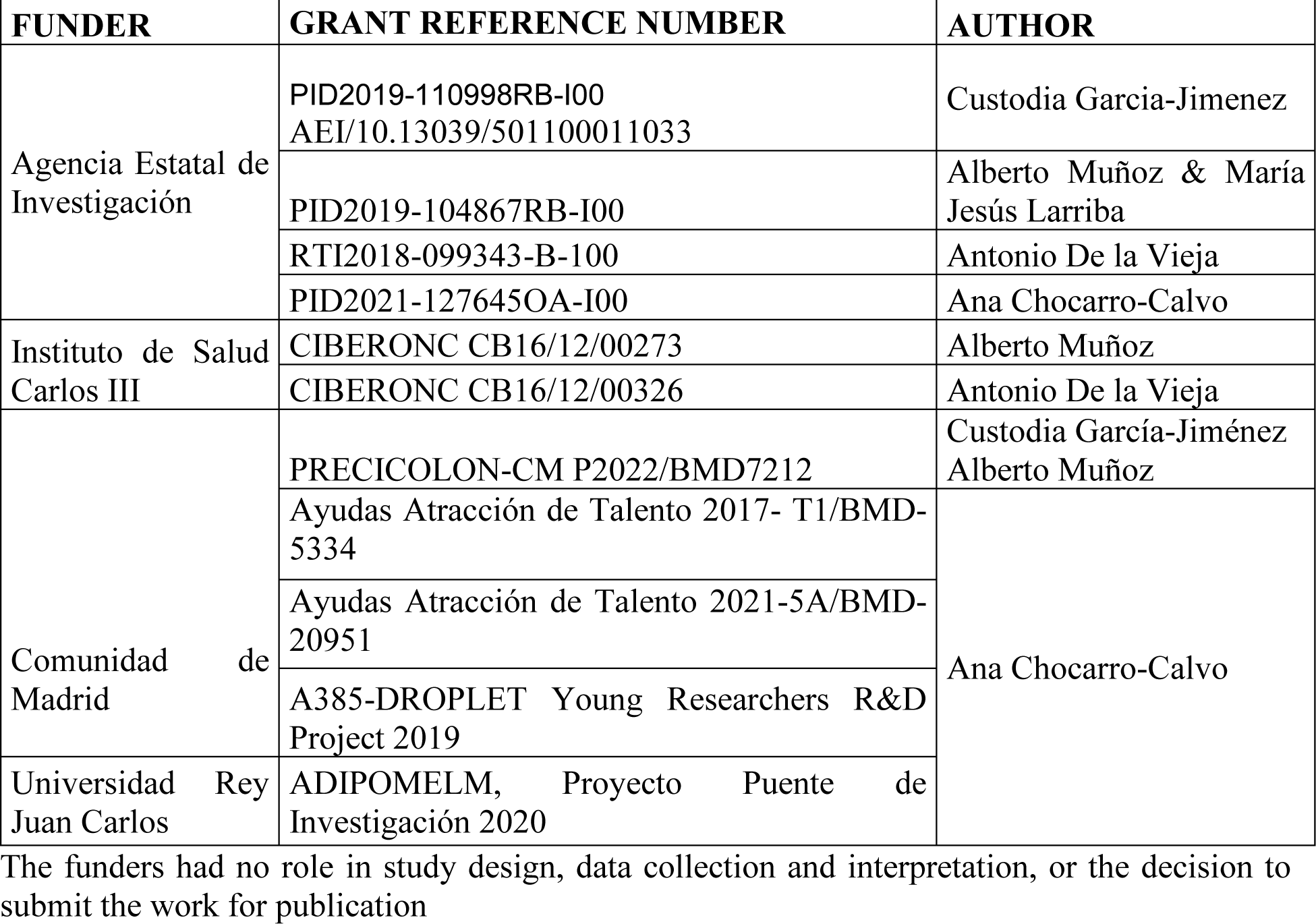

| FUNDER | GRANT REFERENCE NUMBER | AUTHOR |
| --- | --- | --- |
| Agencia Estatal de Investigación | PID2019-110998RB-I00<br>AEI/10.13039/501100011033 | Custodia Garcia-Jimenez |
|  | PID2019-104867RB-I00 | Alberto Muñoz & María Jesús Larriba |
|  | RTI2018-099343-B-100 | Antonio De la Vieja |
|  | PID2021-127645OA-I00 | Ana Chocarro-Calvo |
| Instituto de Salud Carlos III | CIBERONC CB16/12/00273 | Alberto Muñoz |
|  | CIBERONC CB16/12/00326 | Antonio De la Vieja |
| Comunidad de Madrid | PRECICOLON-CM P2022/BMD7212 | Custodia García-Jiménez<br>Alberto Muñoz |
|  | Ayudas Atracción de Talento 2017- T1/BMD-5334 | Ana Chocarro-Calvo |
|  | Ayudas Atracción de Talento 2021-5A/BMD-20951 |  |
|  | A385-DROPLET Young Researchers R&D Project 2019 |  |
| Universidad Rey Juan Carlos | ADIPOMELM, Proyecto Puente de Investigación 2020 |  |

The funders had no role in study design, data collection and interpretation, or the decision to submit the work for publication

## Author contributions

CGJ, JMGM, AM, and MJL conceived the experiments, JMGM, ACC, AB, JMU, MJFA, MCF, JCR performed the experiments. JMGM and ADLV performed the Figs. All authors participated in the discussion. CGJ, AM and MJL wrote the manuscript and provided supervision.

## Ethics

The authors declare that they have no conflict of interest. All procedures followed were in accordance with the ethical standards of the responsible committee on human experimentation (institutional and national) as specified under the paragraph Human samples in the Materials and Methods Section and with the Helsinki Declaration of 1975, as revised in 2000. lnformed consent was obtained from all patients for being included in the study.

## Notes

### Competing Interest Statement

The authors have declared no competing interest.

### Summary of Updates

This version contains slight modifications of the text figures and figure legends to precise and clarify the message answering the referees of eLife.

